# ExoBow: A transgenic strategy to study CD63 exosomes *in vivo*

**DOI:** 10.1101/2021.03.10.434407

**Authors:** Bárbara Adem, Nuno Bastos, Carolina F. Ruivo, Patrícia F. Vieira, Barbara Seidler, José C. Machado, Dieter Saur, Dawen Cai, Sonia A. Melo

**Affiliations:** i3S - Instituto de Investigação e Inovação em Saúde, Universidade do Porto, Portugal; Institute of Molecular Pathology and Immunology of University of Porto, Ipatimup, Portugal; Instituto de Ciências Biomédicas de Abel Salazar, University of Porto, Portugal; Faculty of Medicine, University of Coimbra, Portugal; Medical Clinic and Polyclinic II, Klinikum rechts der Isar, Technical University Munich, Germany; German Cancer Research Center (DKFZ) and German Cancer Consortium (DKTK), Heidelberg, Germany; Department of Pathology, Faculty of Medicine, University of Porto, Portugal; University of Michigan Medical School, Ann Arbor, Michigan, USA

## Abstract

Exosomes are described as central players in a myriad of biological processes. However, the available methodologies to study their function in complex biological systems *in vivo* are still very limited. The biodistribution of endogenously produced exosomes, the ability to trace their spontaneous flow in order to identify the cell types they interact with, remains a major challenge. New tools to identify comprehensive networks of communication established by exosomes originated in distinct cell types *in vivo*, are fundamental for a better understanding of their biology. Here, we describe the development of a genetically engineered mouse model that allows the expression of the mouse CD63 exosomal marker fused with one (monocolor) or up to four fluorescent proteins (multireporter), the ExoBow. The genetic design of the ExoBow transgene allows the conditional expression of the reporters in any tissue/cell-type in an inducible or non-inducible fashion. In addition, communication mediated by CD63 positive (CD63^+^) exosomes can be identified amongst the same tissue/cell types using the multireporter version of the model, in order to map intra-organ/tissue communication. We demonstrate the applicability of the ExoBow transgene in normal physiological conditions and in the context of cancer, using pancreas as a working model. The ExoBow comprises a unique strategy to identify intra- and inter-organ/cell-type communication mediated by CD63^+^ exosomes. We believe this tool will contribute for a better understanding of the complex interactions occurring *in vivo* that underly the biology of exosomes in health and disease.

## INTRODUCTION

Exosomes are small extracellular vesicles (EVs) of endosomal origin, enclosing proteins, lipids, DNA and RNA [1-3]. Through interaction with specific molecular components at the membrane of target cells, or through fusion and delivery of their cargo, exosomes are described to modulate cells locally and at distance [4, 5]. These vesicles have been extensively implied in a plethora of biological processes, including the maintenance of the homeostasis of organs, modulation of the immune system, and pathological conditions such as neurodegenerative diseases and cancer [6-8]. Hence, the need to address the endogenous biodistribution of exosomes and to determine the biological processes they are involved in *in vivo*, is of utmost importance. These studies may corroborate our current knowledge, but also contribute for the discovery of novel features and functions of exosomes, providing new insights in the field. However, the lack of effective tools to determine the whereabouts of endogenous exosomes inside a multicellular organism still undermines the translational potential of most of the data exclusively obtained *in vitro*.

CD63, in addition to CD81 and CD9, is a well-established exosomal marker [1, 9]. Hence, we developed a CD63 multireporter genetically engineered mouse model (GEMM) to determine the spatiotemporal biodistribution of CD63 positive (CD63^+^) exosomes originated in specific organs/cell-types, the ExoBow. The ExoBow transgene can be used both as a monocolor, or a multireporter mouse model. Notwithstanding its innumerous applications, here we demonstrate the use of the ExoBow transgene to study exosomes in normal physiological conditions of the pancreas and in the context of pancreatic cancer.

## RESULTS

### Development of a CD63-XFP genetically engineered mouse model

The transgene (*R26*^*CD63-XFP/*+^) was designed with the purpose to be conditional to any organ/cell-type, and with the possibility to be expressed in an inducible fashion (**Figure 1A**). Briefly, a stop cassette flanked by Frt sites was placed upstream of the CD63 open reading frame (ORF), preventing its expression. The mouse tetraspanin *CD63* is followed by four different fluorescent reporter proteins, mCherry, phiYFP, eGFP and mTFP. The first three reporters are flanked with distinct and incompatible lox recombination sites (*LoxN, Lox2272* and *Lox5171*). Each fluorescent reporter ends with a polyA sequence that enhances expression by stabilizing the transcript and preventing its degradation [10]. Each one of the fluorescent reporters are fused to the C-terminal of the mouse CD63, resulting in a CD63-XFP fusion protein (**Figure 1A**). The transgene was cloned into intron 1 of *ROSA26* locus and is under the control of the *CAG* promoter (CMV enhancer, chicken beta-Actin promoter and rabbit beta-Globin splice acceptor site). To allow the expression of the ExoBow transgene, the flippase (Flp) recombinase excises the stop cassette upstream of the CD63 ORF resulting in the expression of CD63-mCherry fusion protein (the cell is mCherry positive and secretes CD63-mCherry positive exosomes; **Figure 1B**), which represents the CD63-mCherry monocolor mouse model (**Figure 1C**). To achieve the expression of the additional colors, Cre recombinase stochastically selects one of the three pairs of the mutually exclusive lox sites (*LoxN, Lox2272* or *Lox5171*) resulting in cells and respective CD63^+^ exosomes tagged with one of the three additional colors (phiYFP, eGFP and mTFP, respectively; **Figure 1B**). This represents the multireporter mouse model, hereafter referred to as ExoBow (**Figure 1D**). The expected colors of the ExoBow upon the action of Flp and Cre recombinase are mainly phiYFP, eGFP and mTFP. mCherry expression is highly unlikely to occur, although not impossible. For this last event to take place, Flp but not Cre-mediated recombination would have to occur, which is unexpected due to the high efficiency of the Cre recombinase. Further, Cre-mediated recombination can only occur once in each cell, because only one of each of the lox site variants is maintained upon the first Cre recombination event (**Figure 1B**). Recombination efficiency by Flp and Cre recombinases was evaluated in positive embryonic stem cell (ESC) (**Figure 2A** and **2B**, respectively). Recombinant DNA fragments were detected at the expected molecular weight for each recombination site, which validates the recombination efficiency of the ExoBow transgene (**Figure 2A** and **2B**). The ExoBow, on the contrary to the monocolor mouse model, allows the study of communication amongst the same tissue or cell types. In addition, the presence of communication originating from cells and respective CD63^+^ exosomes with distinct colors, provides information on the topographical positioning of the communication routes. Increased frequency of communication taking place in specific routes might be related to close-proximity of the cells. On the other hand, increased frequency in routes of communication between distant cells, and not close ones, can highlight an underlying specificity in that communication avenue.

**Figure 1.**
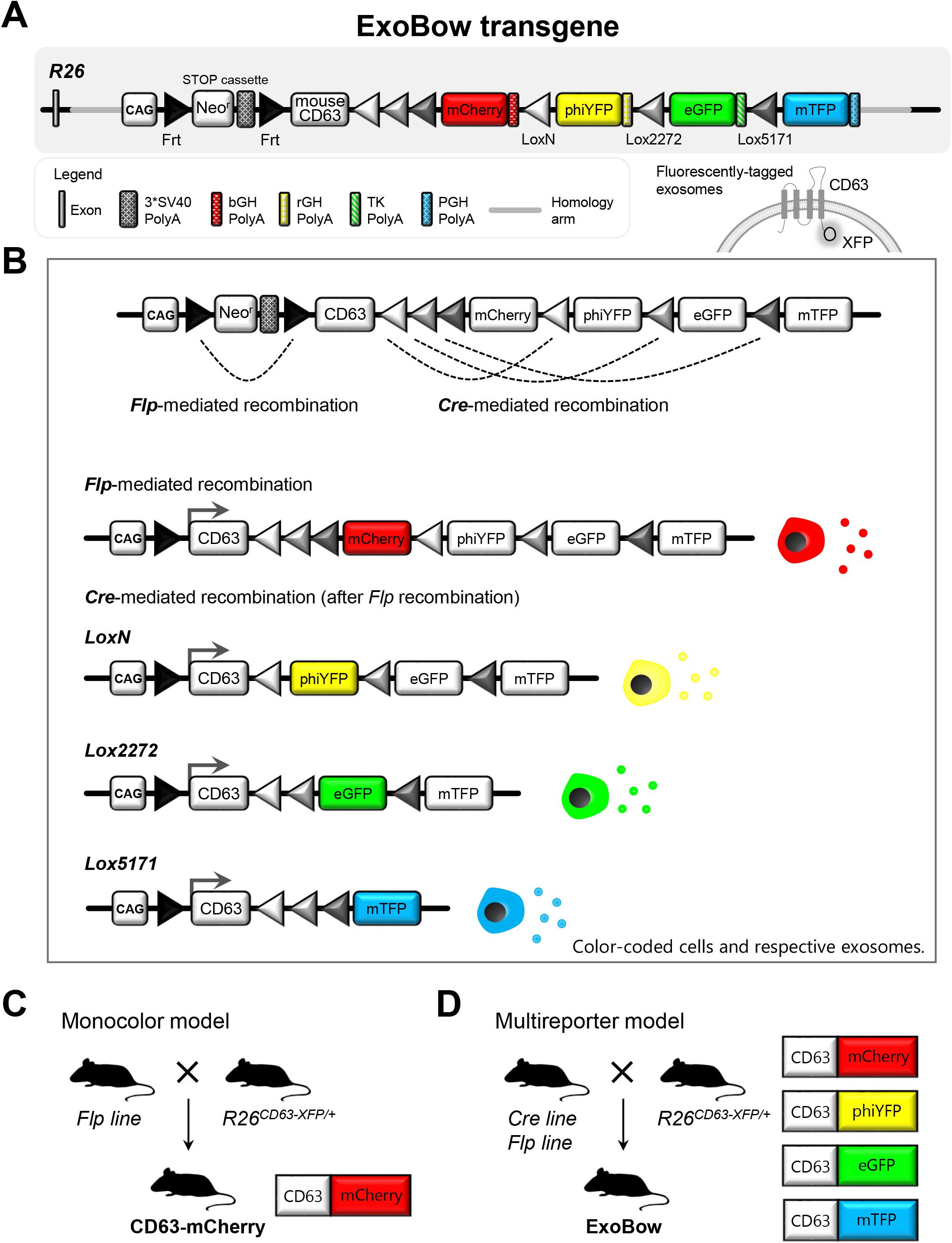
Representation of the ExoBow transgene and recombination strategy. **(A)** The ExoBow construct was inserted in the intron one of *ROSA 26* (*R26*) locus and is under the action of a strong synthetic promotor (CAG). Upstream of the exosomal marker *CD63* gene (mouse) there is a neomycin resistance cassette with a stop-codon flanked by Frt sites that prevents further transcription. Following CD63 there are 4 fluorescent reporters: mCherry, phiYFP, eGFP and mTFP, each with a polyA sequence. Schematic illustration depicting the orientation of the fusion protein CD63-XFP in an exosome. **(B)** When flippase is present it recognizes the Frt sites and excises the stop cassette, allowing CD63-mCherry transcription. Thus, the expressing cell would be mCherry positive and would produce mCherry positive exosomes. Further, Cre-mediated recombination induces the expression of CD63-phiYFP, CD63-eGFP or CD63-mTFP, since the fluorescent reporters are flanked by a different set of lox recombination sites (loxN, lox2272 and lox5171, respectively). The deletions are mutually exclusive preventing further recombination. Hence, cells will have one of the four colors and produce exosomes tagged with the same color, which is maintained upon cell division. **(C)** Schematic of the breeding strategy to obtain the monocolor mouse model CD63-mCherry: cross of a Flp-driven mouse model with the *R26*^*CD63-XFP/*+^ mouse model. **(D)** Schematic of the breeding strategy to obtain the multireporter mouse model ExoBow: cross of a Flp- and Cre-driven mouse models with the *R26*^*CD63-XFP/*+^ mouse model.

**Figure 2.**
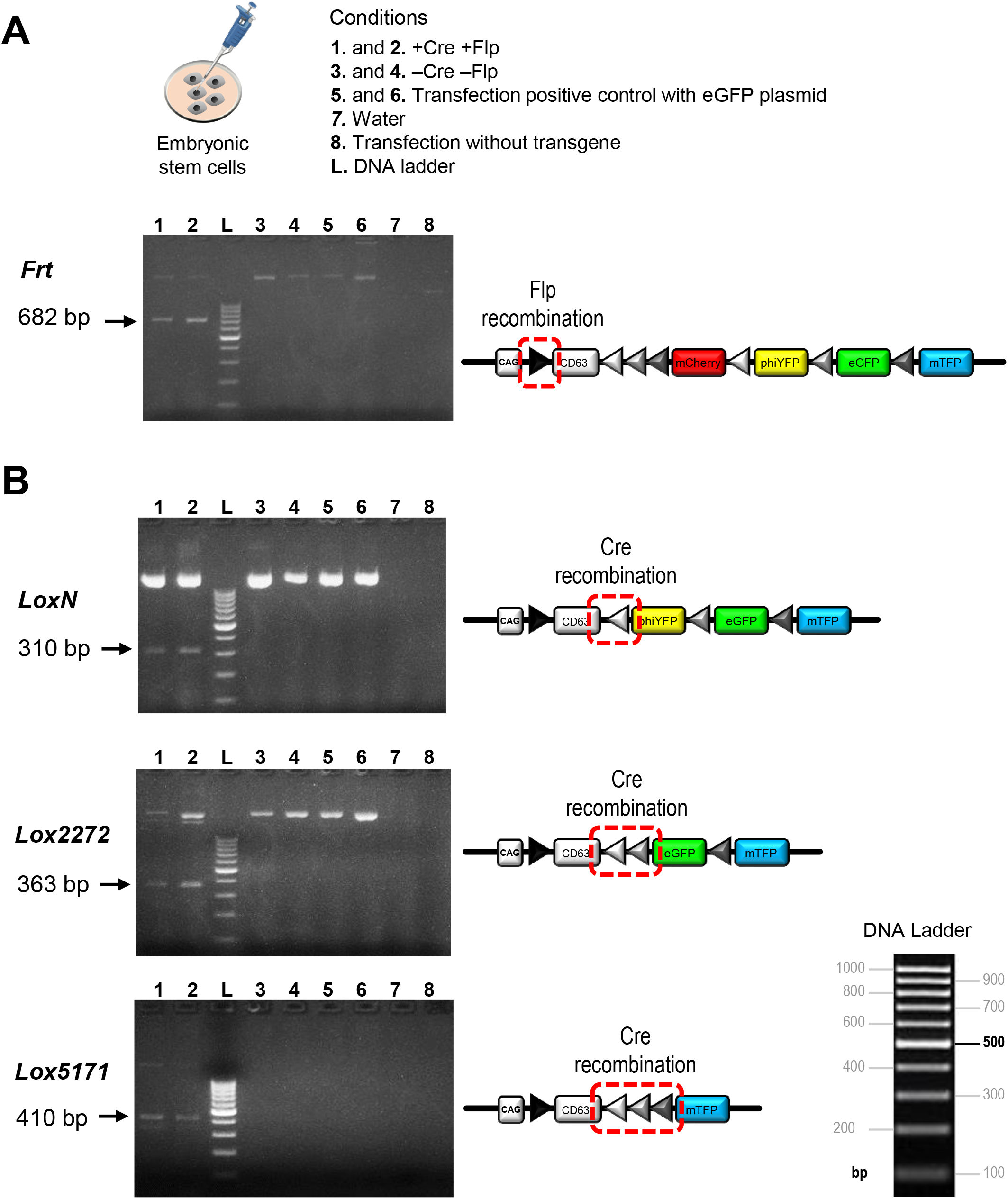
Efficient recombination of the ExoBow transgene in embryonic stem cells mediated by Flp and Cre recombinases. **(A)** Schematic representation of the assessed treatment conditions in embryonic stem cells by PCR. PCR gel depicting Flp-mediated recombination. **(B)** PCR gel depicting Cre-mediated recombination of Lox N sites (upper panel), Lox2272 sites (middle panel) and Lox5171 sites (lower panel). Arrows indicate PCR product in the expected size upon recombination. Control refers to embryonic stem cells transfected with eGFP plasmid (positive control for transfection).

It is important to note that we have inserted only one copy of the transgene in the *ROSA26* locus to assure that, when used in heterozygosity, each cell expresses only mCherry, phiYFP, eGFP or mTFP, and not a combinatorial expression of the different fluorescent reporters. Nonetheless, if the transgene is used in homozygosity, considering the possible combinations, there could be up to 10 colors present. This scenario is assuming that the two fluorescent reporters expressed in one cell and in the produced exosomes co-localize and have the same expression levels. If we assume these conditions, although communication between CD63-XFP expressing cells is virtually impossible to trace, communication from CD63-XFP expressing cells with cells of the microenvironment, or with cells at other organs can be possible. Analysis of the homozygous version can be very challenging, thus in here we have used the multireporter model in heterozygosity (**Supplementary Figure 1**, gel of heterozygous *versus* homozygous R26^CD63-XFP^ mice).

### CD63-XFP proteins co-localize with endogenous CD63 and efficiently tag CD63^+^ exosomes

To assess if CD63-XFP proteins co-localize in the same cellular compartments as the endogenous CD63, the exact sequence of each mouse CD63-XFP from the ExoBow transgene was cloned into the pRP[Exp]-Puro-CAG backbone vector and used to transfect a human pancreatic cancer cell line (BxPC-3; **Figure 3A**). BxPC-3 cells were separately transfected with the CD63-mCherry, CD63-phiYFP, CD63-eGFP and CD63- mTFP plasmids to obtain stable clones expressing only one of the CD63-XFP proteins. We found that the endogenous (human) CD63 protein presents a speckle-like pattern with accumulation in proximity to the nuclei, suggestive of endosomal location, as expected (**Figure 3B**) [11]. In addition, the expression of the CD63-XFP proteins was identical to the endogenous CD63 of the parental cell line (**Figure 3C**). To further confirm that each mouse CD63-XFP protein co-localizes with the endogenous human CD63, we have used anti-XFP antibodies (anti-mCherry, anti-phiYFP and anti-mTFP developed by Cai Laboratory derived from the Brainbow tools [12]), and a specific anti-human CD63 antibody (to detect endogenous human CD63). We found that the mouse CD63-XFP proteins co-localize with the endogenous human CD63, confirming that CD63 fusion with reporter proteins does not interfere with their cellular location (**Figure 3D**). Anti-human CD63 antibody specificity was confirmed by immunofluorescence using a mouse cell line (KPC; **Supplementary Figure 2A**). Next, we evaluated the presence of CD63-XFP color-coded exosomes derived from each clone. Extracellular vesicles were isolated by ultracentrifugation followed by a continuous sucrose gradient (**Figure 3E**). Protein isolated from fractions corresponding to distinct densities were analyzed by western-blot using anti-XFP antibodies, demonstrating that the distinct CD63-XFP clones secrete color-coded exosomes (**Figure 3E**). Protein fractions derived from the sucrose gradient of the vesicles from the parental BxPC-3 cell line were immunoblotted against mCherry as control (**Supplementary Figure 2B**).

**Figure 3.**
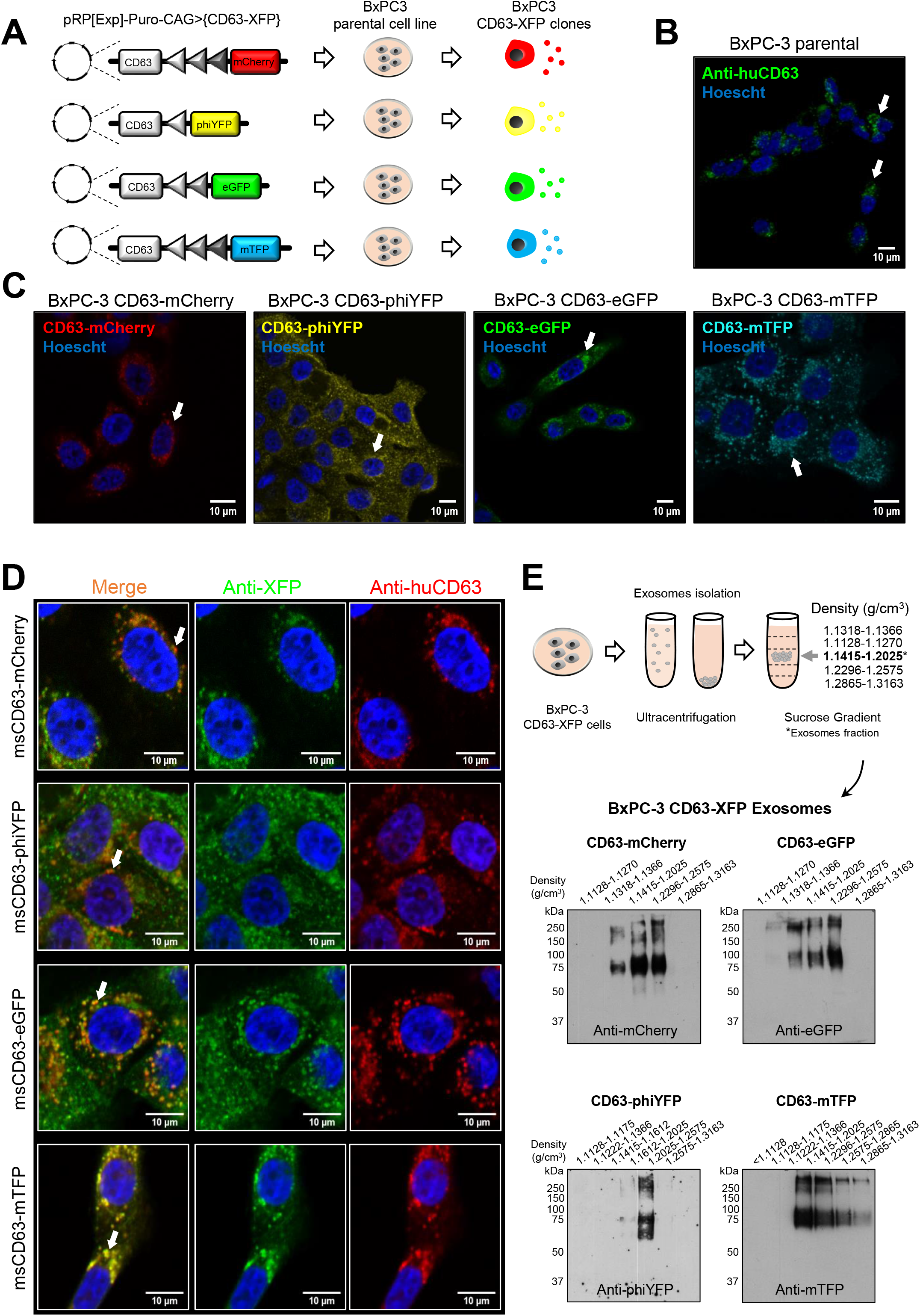
CD63-XFP PDAC cell lines produce color-coded exosomes. **(A)** Each CD63-XFP (CD63 of mouse origin) sequence that result from the Flp and Cre mediated recombinations of the ExoBow transgene was cloned into the pRP[Exp]-Puro-CAG> plasmid and transfected into the human pancreatic ductal adenocarcinoma cell line BxPC-3. **(B)** Confocal microscopy image of maximum projection of BxPC-3 parental cells immunostained against anti-human CD63 (endogenous CD63 of BxPC-3 parental cell line). Arrows depict the perinuclear accumulation of CD63. **(C)** Confocal microscopy images of the fluorescent endogenous levels of each BxPC-3 CD63-XFP clone: CD63- mCherry (red), CD63-phiYFP (yellow), CD63-eGFP (green) and CD63-mTFP (cyan), using anti-XFP antibodies. The nuclei counterstained with Hoescht are represented in blue. Arrows depict the perinuclear accumulation of CD63-XFP. **(D)** Confocal microscopy images of BxPC-3 CD63-XFP cells immunostained with respective anti-XFP antibody (green) and the anti-human CD63 antibody (red) to determine cellular location. Nuclei counterstained with Hoescht are represented in blue. Arrows indicate examples of co- localization sites. **(E)** Schematic representation of exosomes isolated from each BxPC3 CD63-XFP clone by continuous sucrose gradient protocol. Individual 1 mL fractions were collected and after ultracentrifugation were loaded on gels for electrophoresis. Exosomes are located in fractions around 1.1415 to 1.2025 g/cm^3^ density. Anti-mCherry, anti-eGFP, anti-phiYFP and anti-mTFP antibodies were used in the respective CD63- mCherry, CD63-eGFP, CD63-phiYFP and CD63-mTFP western-blots.

### Panc-CD63-mCherry and Panc-ExoBow genetically engineered mouse models

To achieve a pancreas-specific expression of the ExoBow transgene, we crossed this model with the *Pdx1-Flp* to generate the Panc-CD63-mCherry (monocolor model) [13]. In this model, the recombinase Flp is under the control of *Pdx1*, a pancreas-specific promoter, leading to the expression of CD63-mCherry (**Figure 4A**). Hence, the cells of the pancreas are CD63-mCherry positive (CD63-mCherry^+^) and produce CD63- mCherry^+^ exosomes. Using IVIS Lumina system we observed a strong fluorescent signal in the pancreas of the Panc-CD63-mCherry mice (*Pdx1-Flp*; *R26*^*CD63-XFP/*+^), while the control (*R26*^*CD63-XFP/*+^) did not show mCherry signal when using a 535 excitation laser and a DsRed emission filter (**Figure 4B**). In addition, flow cytometry analysis of the pancreas of *WT* and Panc-CD63-mCherry mice showed CD63-mCherry^+^ cells only in the later, demonstrating an efficient removal of the STOP cassette by Flp recombinase (**Figure 4C**). Immunofluorescence analysis of the pancreas of Panc-CD63-mCherry mice also demonstrate the efficient recombination of the transgene (**Figure 4D**). In addition, the expression of the CD63-mCherry fusion protein is observed in the cells of the exocrine as well as endocrine pancreas (islet of Langerhans) (**Figure 4D**). Pancreas cells show a CD63-mCherry speckle-like pattern of expression suggestive of endosomal localization (**Figure 4E**). Furthermore, EVs isolated from the pancreas of Panc-CD63-mCherry mice are mCherry^+^ on the contrary to the ones isolated from *WT* mice (**Figure 4F**). Notably, the small EVs are significantly more enriched in CD63-mCherry than the larger EVs subset (**Figure 4F**). Altogether, demonstrating that pancreas cells of Panc-CD63- mCherry mice secretes CD63-mCherry^+^ exosomes.

**Figure 4.**
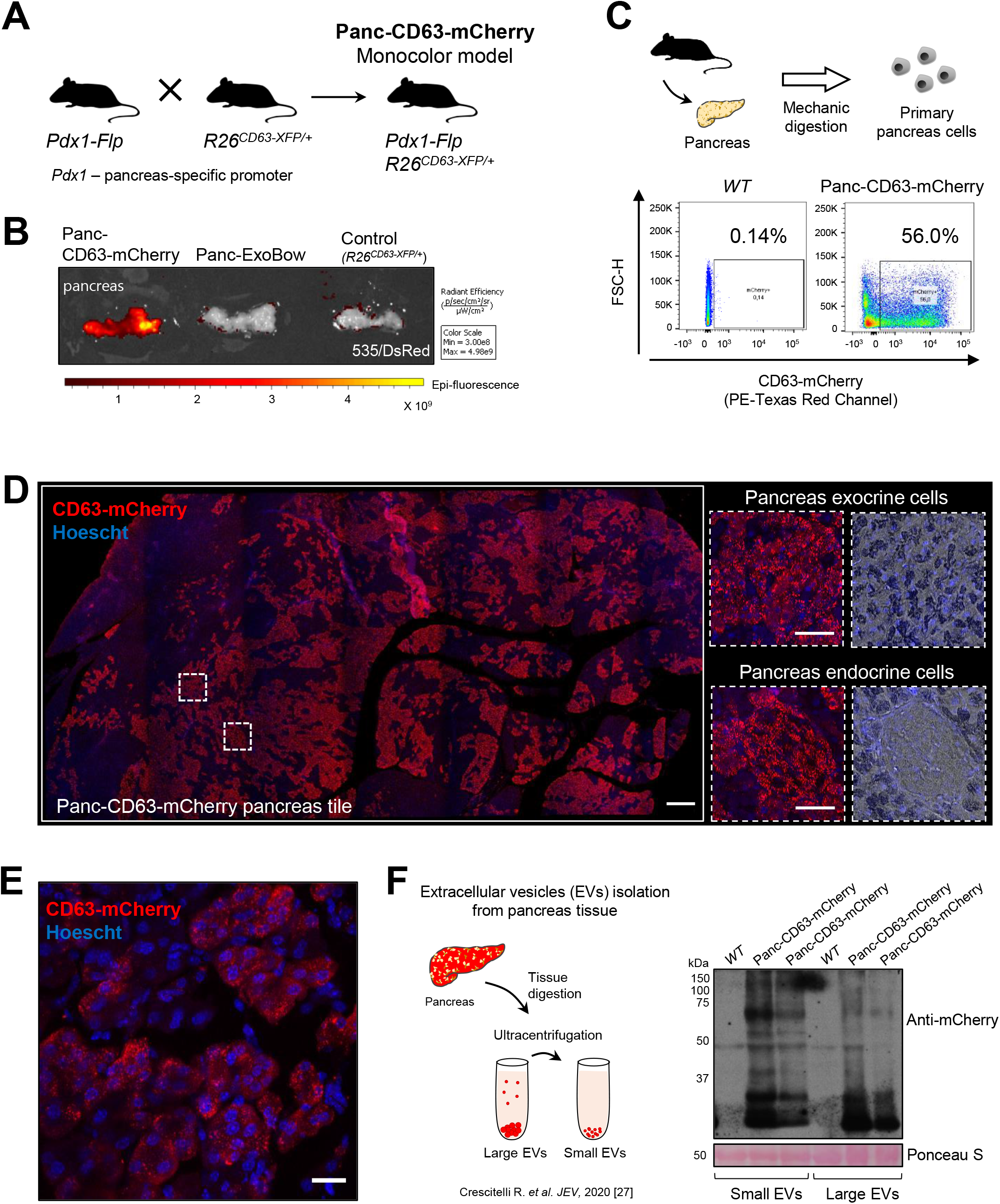
Panc-CD63-mCherry portrays efficient ExoBow transgene recombination and production of color-coded exosomes. **(A)** Schematic of the breeding strategy to obtain the monocolor mouse model, Panc-CD63- mCherry. Flp recombinase is under the control of a pancreas-specific promoter, *Pdx1*, allowing a conditional expression of the ExoBow transgene in pancreas cells **(B)** Pancreas imaging using IVIS Lumina System illustrating CD63-mCherry expression (same image/pancreas as in Figure 5B, but here we are using a 535 excitation laser and DsRed emission filter). Pancreas of a monocolor mouse model Panc-CD63-mCherry (left), multireporter mouse model Panc-ExoBow (middle) and control (*R26*^*CD63-XFP/*+^, no recombinases) (right). **(C)** Representative flow cytometry analysis of CD63-mCherry positive cells in a pancreas of a Panc-CD63-mCherry mouse (right) and the pancreas of a *wild-type* (*WT*) mouse (control) (left). **(D)** Confocal microscopy images of a tile scan maximum projection depicting CD63-mCherry positive cells in the pancreas of a Panc- CD63-mCherry mouse. Immunofluorescence against mCherry (red) and nuclei counterstained with Hoescht (blue). Insets of pancreas exocrine and endocrine cells with respective brightfield images merged with Hoescht. Scale bar 200μm. **(E)** Confocal microscopy images of a maximum projection of a pancreas section depicting exocrine CD63-mCherry positive cells in the Panc-CD63-mCherry model. Immunofluorescence against mCherry (red) and nuclei counterstained with Hoescht (blue). Scale bar 20μm. **(F)** Schematic representation of the isolation of interstitial extracellular vesicles (EVs) from the pancreas tissue by ultracentrifugation according to Crescitelli *et al*. [27]. Anti-mCherry western-blot in small and large EVs fractions isolated from pancreas tissue of wild-type (*WT*, control) or Panc-CD63-mCherry mice. The CD63-mCherry fusion protein appears at the expected molecular weight between 50 and 75 kDa. Band around 27 kDa identify the mCherry alone protein as a result of the harsh tissue digestion.

In order to obtain the multireporter version of the ExoBow transgene, in addition to Flp driven recombination, Cre recombinase action is also required. Thus, we crossed the ExoBow with the *Pdx1-Flp* and *Pdx1-Cre* alleles to generate a multireporter model conditional to the pancreas, the Panc-ExoBow (**Figure 5A**). Using IVIS Lumina system with a 465-excitation laser and a GFP emission filter (it detects eGFP, phiYFP and mTFP), we observed a strong fluorescent signal in the pancreas of the Panc-ExoBow mice (*Pdx1-Flp*; *Pdx1-Cre*; *R26*^*CD63-XFP*^) while no signal was observed in the control (*R26*^*CD63-XFP/*+^) and the CD63-mCherry mice (*Pdx1-Flp*; *R26*^*CD63-XFP/*+^; **Figure 5B**). This demonstrates the successful recombination of the ExoBow transgene. The expression of the different CD63-XFP proteins by pancreas cells was confirmed by immunofluorescence analysis (**Figure 5C**).

**Figure 5.**
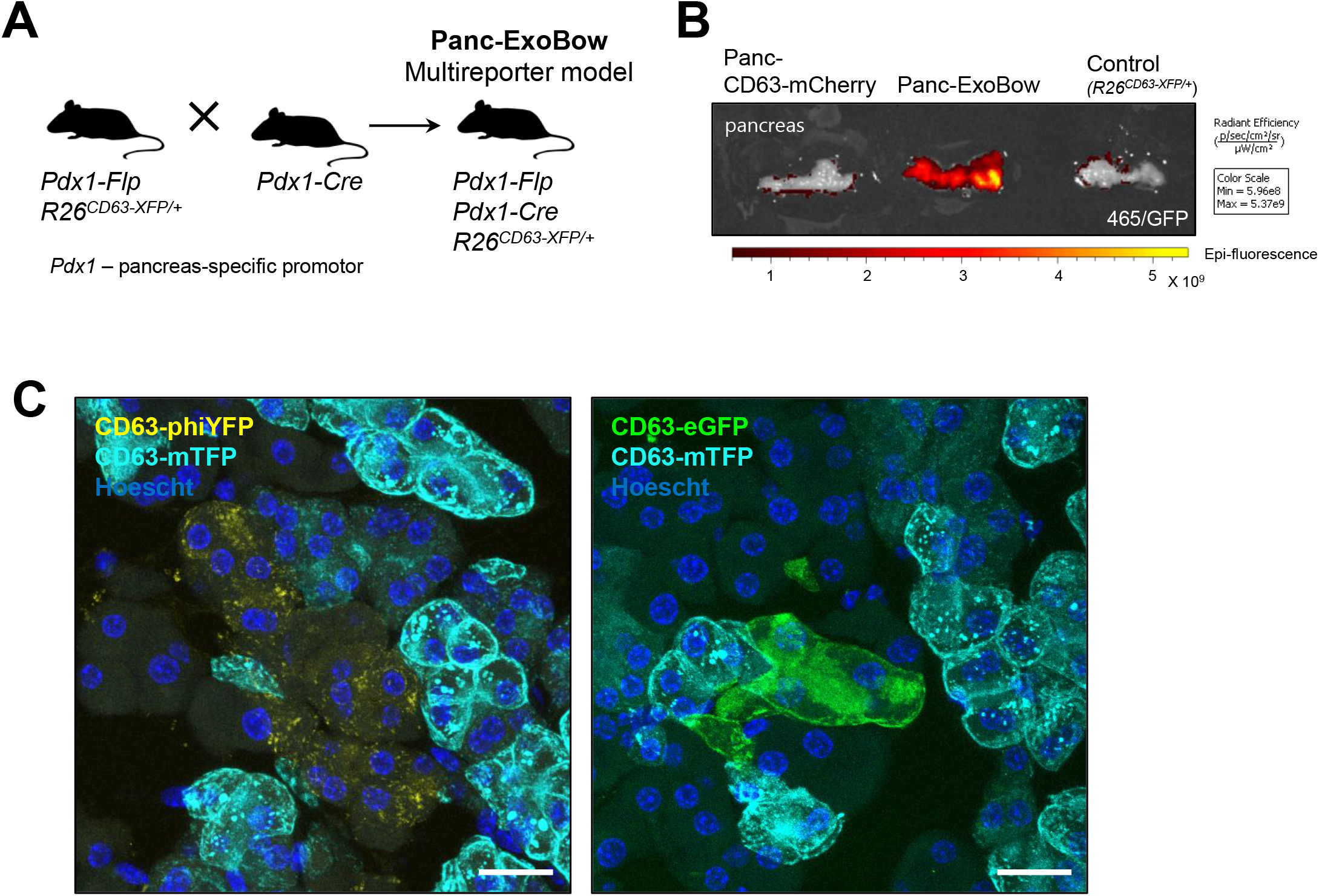
Panc-ExoBow exhibits efficient ExoBow transgene recombination and color-coded CD63-XFP cells. **(A)** Schematic representation of the breeding strategy to obtain the multireporter mouse model, Panc-ExoBow. Flp and Cre recombinases are under the control of a pancreas-specific promoter, *Pdx1*, allowing a conditional expression of the ExoBow transgene in pancreas cells. **(B)** Pancreas imaging using IVIS Lumina System illustrating CD63-eGFP, CD63-phiYFP, and CD63-mTFP (same image/pancreas as in Figure 4B but here we are using a 465 excitation laser and GFP emission filter). Pancreas of a monocolor mouse model Panc-CD63 mCherry (left), multireporter mouse model Panc-ExoBow (middle) and control (*R26*^*CD63-XFP/*+^, no recombinases) (right). **(C)** Confocal images of a maximum projection of a pancreas section depicting CD63-mTFP (cyan), CD63-phiYFP (yellow) and CD63-eGFP (green) positive cells in the Panc-ExoBow model. Immunofluorescence for mTFP, phiYFP and eGFP, and nuclei were counterstained with Hoescht (blue). Scale bar 20μm.

### Pancreatic ductal adenocarcinoma ExoBow genetically engineered mouse models (PDAC-ExoBow)

The ability of the ExoBow transgene to color pancreatic cancer cells was evaluated using two distinct pancreatic ductal adenocarcinoma (PDAC) GEMMs: Flp-driven KPF model (*Pdx1-Flp*; *FSF-Kras*^*G12D/*+^; *Trp53*^*Frt/*+^) and the Cre-driven KPC model (*Pdx1-Cre*; *LSL- Kras*^*G12D/*+^; *LSL-Trp53*^*R172H/*+^). The KPF presents the conditional expression of the Kras^G12D^ mutation and heterozygous loss of the *Trp53* allele driven by conditional *Flp* recombination in pancreas cells through the *Pdx1* promoter, leading to the spontaneous development of PDAC [13]. Crossing the KPF with the ExoBow allele results in Flp-mediated recombination of the ExoBow transgene allowing the expression of CD63- mCherry in cancer cells, corresponding to the PDAC monocolor mouse model, KPF CD63-mCherry (*Pdx1-Flp; R26*^*CD63-XFP-/-*^; *FSF-Kras*^*G12D/*+^; *Trp53*^*Frt/*+^) (**Figure 6A**). IVIS Lumina System analysis of the pancreas of KPF CD63-mCherry mice showed an intense and specific signal in comparison with a pancreas from a control animal (no ExoBow transgene), demonstrating the successful recombination of the ExoBow transgene (**Figure 6B**). Immunofluorescence analysis of the pancreas also demonstrates a significant percentage of the pancreas expressing CD63-mCherry^+^ cells (**Figure 6C**). PDAC tumors are characterized by an extensive desmoplastic reaction, and the tumors are populated of cancer associated fibroblasts (CAFs) [14]. Communication between cancer cells and CAFs, mediated or not by vesicles, has been extensively described [5, 15]. As a proof of concept, we demonstrate in tumors of the KPF CD63-mCherry model that cancer CD63^+^ exosomes enter spontaneously in CAFs (**Figure 6D**).

**Figure 6.**
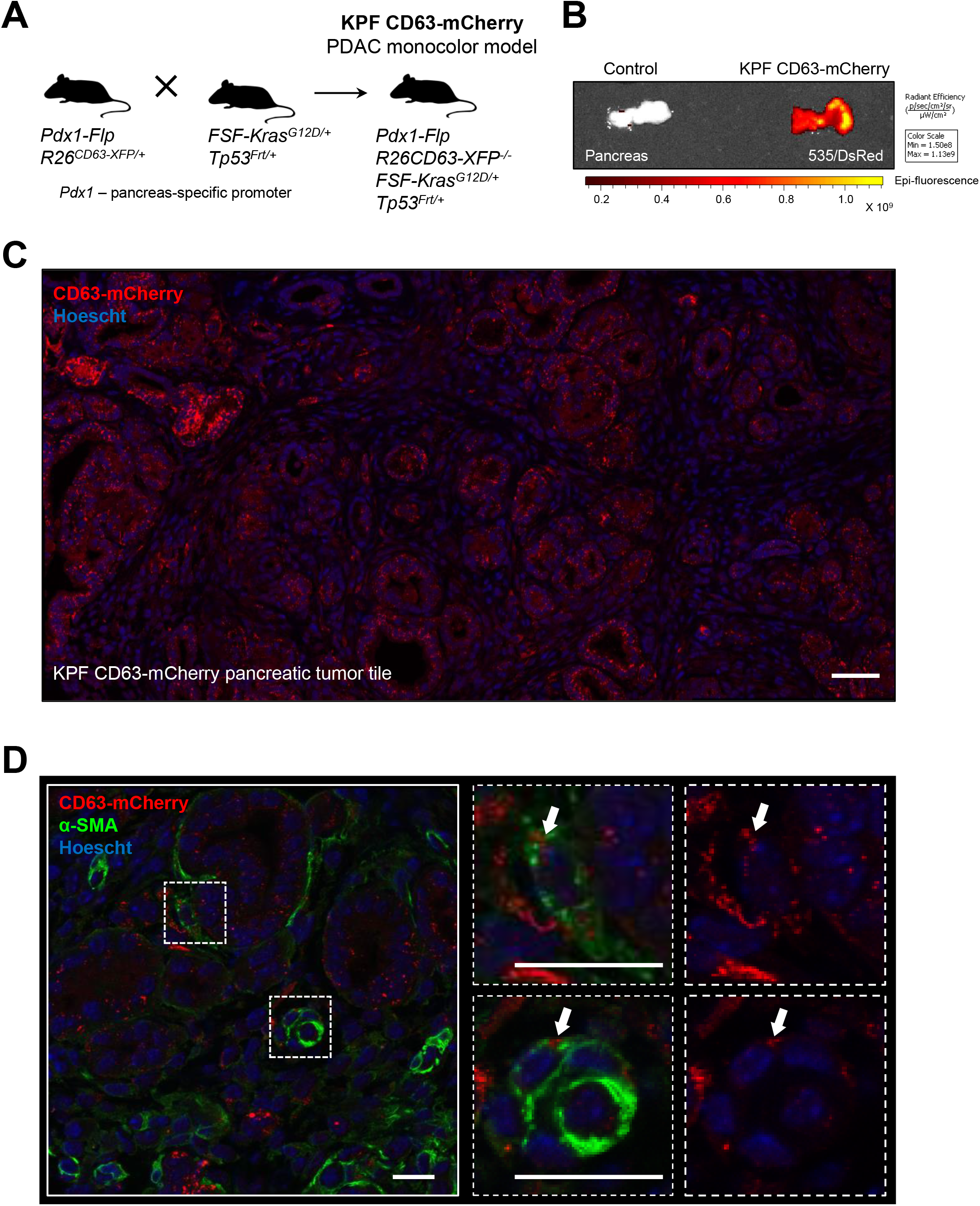
KPF CD63-mCherry exhibits efficient ExoBow transgene recombination and communication with other cells of the tumor is detected. **(A)** Schematic representation of the breeding strategy to obtain the PDAC monocolor mouse model, KPF CD63-mCherry. Flp recombinase is under the control of a pancreas-specific promoter, *Pdx1*, allowing the expression of the *Kras*^*G12D*^ mutation, loss of the *Trp53* allele and expression of the ExoBow transgene specific to pancreatic cancer cells. **(B)** Pancreas imaging using IVIS Lumina System with 535 excitation laser and DsRed emission filter. Pancreas of a control mouse (no ExoBow transgene) (left) and of a monocolor PDAC mouse model KPF CD63-mCherry (right). **(C)** Confocal microscopy images of a tile scan depicting CD63-mCherry positive cancer cells in the pancreas of a KPF CD63-mCherry mouse. Immunofluorescence against mCherry (red) and nuclei were counterstained with Hoescht (blue). Scale bar 50 μm. **(D)** Confocal microscopy image of immunostaining anti-mCherry (red) and anti-alpha-smooth muscle actin (α- SMA) labeling CAFs (green), and nuclei counterstained with Hoescht (blue). Arrows represent examples of CD63-mCherry positive exosomes in CAFs. Scale Bar 20μm.

Next, we have crossed the Cre-driven KPC model with the ExoBow. KPC mice spontaneously develop PDAC with full penetrance and reliably recapitulate the clinical and histopathological features of the human disease [16, 17]. These mice harbor the *Kras*^*G12D*^ and *Trp53*^*R172H*^ mutations conditionally expressed in pancreas cells through the action of Cre recombinase under the control of the *Pdx*1 promoter [16]. Thus, crossing the KPC with the ExoBow and the *Pdx1-Flp* alleles we generated the KPC-ExoBow (*Pdx1-Flp; Pdx1-Cre; R26*^*CD63-XFP/*+^; *LSL-Kras*^*G12D/*+^; *LSL-Trp53*^*R172H/*+^) that represents the multireporter version of the PDAC mouse model (**Figure 7A**). In the KPC-ExoBow, Cre recombinase stochastically choses one of the three possible lox sites in the ExoBow transgene, and Flp recombination excises the stop cassette upstream of the CD63-XFP ORF. In this way, each cancer cell expresses one of the three possible fusion proteins (CD63-phiYFP, CD63-eGFP and CD63-mTFP) and secretes the respective CD63-XFP^+^ exosomes. In a similar fashion to what we observed in the KPF CD63-mCherry, we found that cancer cells are CD63-XFP color-coded (**Figure 7B**) and secrete color-coded CD63^+^ exosomes that enter spontaneously in CAFs (**Figure 7C**).

**Figure 7.**
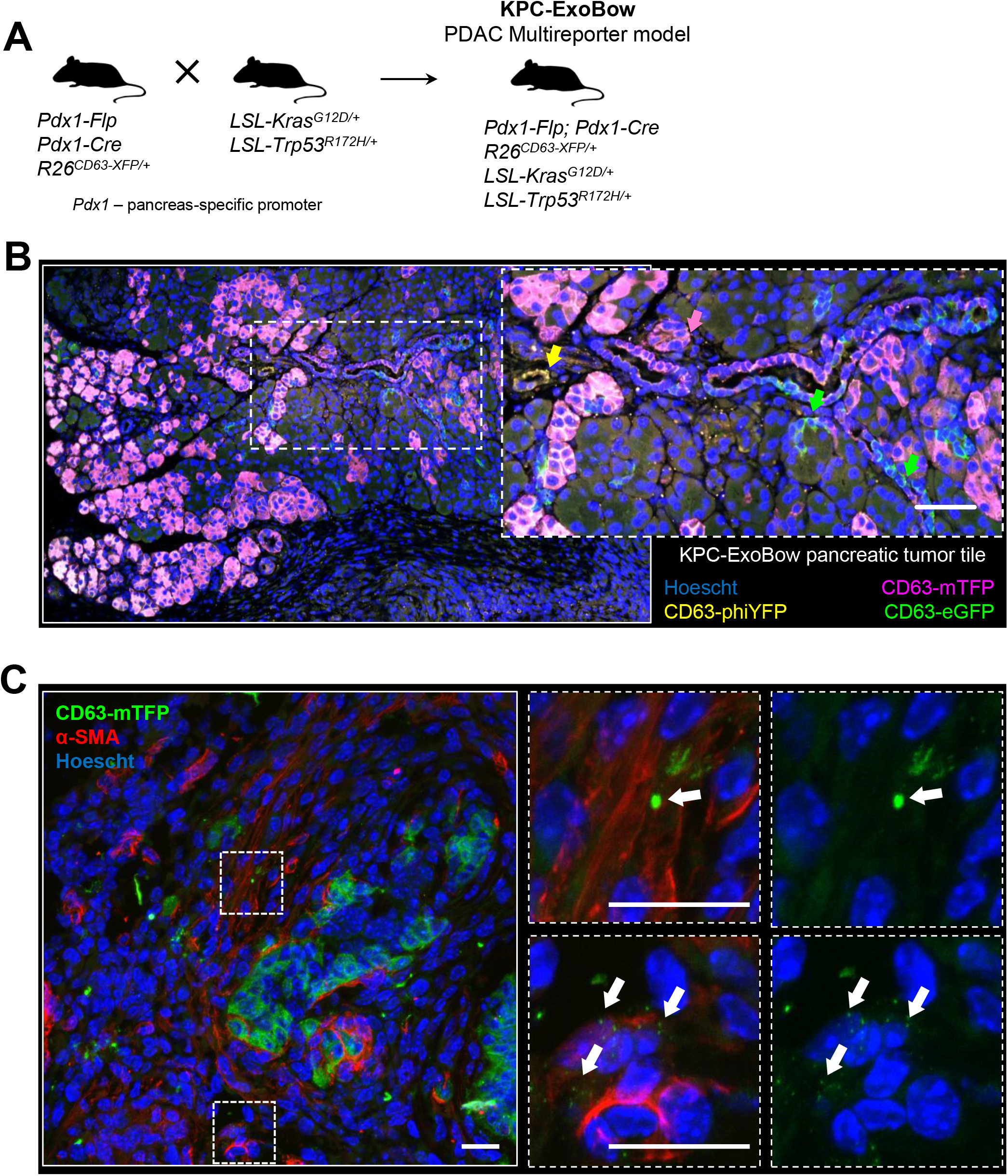
KPC-ExoBow exhibits efficient ExoBow transgene recombination and communication with other cells of the tumor is detected. **(A)** Schematic representation of the breeding strategy to obtain the PDAC multireporter mouse model, KPC-ExoBow. Flp and Cre recombinases are under the control of a pancreas-specific promoter, *Pdx1*, allowing the expression of the KRAS^G12D^ mutation, Trp53^R172H^ mutation and expression of ExoBow transgene specific to pancreatic cancer cells. **(B)** Confocal microscopy images of a tile scan depicting CD63-mTFP, CD63-phiYFP and CD63-eGFP positive cells in the pancreas of a KPC-ExoBow mouse. Scale Bar 200μm. Digital zoom- in to highlight the expression of the different CD63-XFP proteins in pancreatic cancer cells. Scale bar 50μm. Immunofluorescence against mTFP (magenta), phiYFP (yellow) and eGFP (green), and nuclei counterstained with Hoescht (blue). **(C)** Confocal microscopy image of immunostaining against mTFP (green) and alpha-smooth muscle actin (α-SMA) labeling CAFs (red), and nuclei counterstained with Hoescht (blue). Arrows represent examples of CD63-mTFP positive exosomes in CAFs. Scale bar 20μm

## DISCUSSION

The realization of the importance of intercellular communication mediated by EVs in distinct biological processes in health and disease in the past years, has spawn the efforts spent in developing new methodologies and standardization of protocols to improve the biological significance of our findings. Today, we believe that exosomes are important mediators of communication, with roles that span from maintenance of normal homeostasis to several pathological processes. Nonetheless, most of the current knowledge on exosomes biology is supported by *in vitro* experiments or *in vivo* models that poorly mimic physiological conditions. Until recently, *in vivo* studies were limited to the use of exosomes labelled with dyes, or tagged exosomes derived from genetically engineered cells foreigner to the host organism [18]. These approaches present well described limitations, reflected in non-concordant results when we vary the route of administration, the quantity of exosomes and method of isolation used, their source, the use of immunodeficient *versus* immunocompetent mice, amongst others [18, 19]. These drawbacks hamper a clear biological interpretation of the findings.

Recently, there has been a significant investment in the development of new models that enable the study of the biology of exosomes in a context closer to the standard biological system [20-25]. Distinct mouse models that allow the conditional expression of a tetraspanin fused with a reporter have been described, including CD9-GFP [22], CD63- emGFP [23], CD63-NanoLuc [24], and human CD63-GFP [25].

The ExoBow, in addition to the so far developed exosomes-reporter mouse models, opens up the possibility to address distinct biological questions that cannot be answered with any of the existing models. The multireporter version enables the expression of up to 4 fluorescent reporters in heterozygosity. This characteristic allows to interrogate communication within cells of the same cell lineage (e.g. intratumor communication), investigate the topographical positioning of the communication routes in place and understand if cell-cell proximity is a factor that dictates the route and/or frequency of communication. The homozygous version can have up to 10 colors, nonetheless this can be an imaging challenge. In addition, using the same transgene we can also have a conditional monocolor mouse (CD63-mCherry), which confers our transgenic strategy a level of plasticity unforeseen in any of the available models to date.

Despite the advancements, no model is perfect. Cells are described to produce heterogenous populations of extracellular vesicles [9]. Since it was not yet discovered a consensus marker that includes all these populations (most likely there is no such thing as a ubiquitous marker), the current models are limited to the study of one population. Furthermore, the most fit tetraspanin of choice may vary between tissue/cell-type or a pathological context. Finally, the potential effects of overexpressing tetraspanins on mice/disease development, exosomes production, release, distribution, and function cannot be overlooked. The collective findings from different models in the same biological setting will provide new valuable information to so many questions that remain elusive.

## MATERIAL AND METHODS

### Cell Culture

We have used a human and a mouse pancreatic ductal adenocarcinoma cell line, BxPC- 3 (ATCC Cat# CRL-1687) and KPC (Ximbio Cat# 153474), respectively. BxPC-3 and KPC cells were routinely mycoplasma tested. STR profiling was performed in the BxPC- 3 cell line. BxPC-3 cells were cultured in RPMI-1640 medium (Gibco) supplemented with 10% (v/v) fetal bovine serum (FBS, Gibco), 100 U/mL penicillin and 100 μg/mL streptomycin (Gibco), and KPC cells were cultured in DMEM medium (Gibco) supplemented with 10% (v/v) fetal bovine serum (FBS, Gibco), 100 U/mL penicillin and 100 μg/mL streptomycin (Gibco). All stable clones derived from the parental BxPC-3 cell line were cultured in the same conditions. The cells were maintained at 37°C in a humidified chamber with 5% CO^2^.

### Cell line transfection

BxPC-3 cells were separately transfected with the CD63-XFP vectors: CD63-mCherry, CD63-phiYFP, CD63-eGFP and CD63-mTFP that were cloned into pRP[Exp]-Puro-CAG plasmid backbone in collaboration with Vector Builder Inc. Reverse transfection of each plasmid (2.5 µg DNA/1.5 × 10^5^ cells) was performed using Invitrogen Lipofectamine® 2000 (Thermo Fisher Scientific) according to the manufacturer’s instructions. To obtain stable clones, puromycin (1μg/mL Sigma-Aldrich P8833) selection and fluorescence activated cell sorting (FACS) based on the expression of fluorescent proteins was performed.

### Exosomes isolation from cell culture medium and sucrose gradient

BxPC-3 cells were cultured in exosomes-free medium (RPMI-1640 medium (Gibco) supplemented with 10% FBS (Gibco) depleted of exosomes by ultracentrifugation overnight at 100 000g, and 100 U/mL penicillin and 100 μg/mL streptomycin (Gibco). After 72 hours, the medium was collected and centrifuged at 2500g for 10 min followed by a 5 min centrifugation at 4000g. Subsequently, the medium was filtered through a 0.2 μm filter (GE Healthcare Whatman™) directly to an ultra-clear centrifuge tube (Beckman Coulter®). The samples were centrifuged overnight at 100 000g, 4°C using the Optima™ L-80 XP ultracentrifuge, Beckman Coulter. The supernatant was carefully removed, and the isolates were subjected to a continuous sucrose gradient (0.25 – 2 M) as described in [26]. Briefly, the pellet was resuspended in 2 mL of HEPES/Sucrose stock solution (HEPES 20 mM/protease-free sucrose 2.5M, pH 7.4) and transferred to an ultra-clear centrifuge tube. A continuous sucrose solution (2 M to 0.25 M) was dispensed into the ultracentrifuge tube containing the exosomes suspension. Samples were centrifuged overnight (>14 hours) at 210 000g, 4°C. After ultracentrifugation, 1 mL of gradient fractions, from top to bottom, were collected. 50μL of each fraction was used to measure the refractive index in a refractometer. Each fraction was individually placed in an ultra-clear centrifuge tube, diluted in NaCl 0.9% and centrifuged at 100 000g for 2 hours, at 4°C. The subsequent pellet was resuspended in 30μL 2.5% SDS/8 M Urea for protein extraction and incubated for 30 min on ice, followed by a 30 min centrifugation at 17 000g, 4°C. The supernatant was stored at −20°C.

### Production of CAG/mouseCD63-XFP genetically engineered mice using embryonic stem cells

R26^CD63-XFP/+^ allele was developed in collaboration with Cyagen. Briefly, the “CD63- cDNA-LoxN-Lox2272-Lox5171-mCherry-bGHPolyA-LoxN-phiYFP-rBGPolyA-Lox2272- GFP-TKPolyA-Lox5171-mTFP-PGKPolyA” transgene was cloned into intron 1 of ROSA26 and the “CAG-Frt-Stop (Neo cassette with 3*SV40 PolyA)-Frt” was placed upstream of the transgene. The model was generated by homologous recombination in embryonic stem cells.

The diphtheria toxin (DTA) cassette, the ROSA26-homology arms, the CAG promoter and the bGH polyadenylation have been cloned into backbone as prepared. To engineer the targeting vector, the components of neo-3*SV40pA, mcherry-bGHpA, phiYFP- rBGpA, EGFP-TKpA, and mTFP-PGKpA were generated by PCR using gene synthesis product as template. PCR primers were designed to share 15-20 bases of homology with the sequence at the end of the linearized backbone vector.

Mouse genomic fragments containing homology arms (HAs) and conditional knock-in region were amplified from BAC clone (RP23-244D9) by using high fidelity Taq. Next, the fragments were sequentially assembled into a targeting vector together with recombination sites and selection markers. Each individual cloning step was extensively validated through restriction analysis and partial sequencing.

In detail, the neo-3*SV40pA cassette was cloned into Basic Vector by In-Fusion Enzymes, and the Basic Vector came from digested products of NsiI/XhoI. The correct plasmid was named Cd63-Step1. The Cd63-cDNA-3*loxp (NCBI Reference Sequence: NM_007653.3; Ensembl: ENSMUSG00000025351; Transcript: Cd63-201 ENSMUST00000026407.7) were cloned into Cd63-Step1 by In-Fusion Enzymes, and the Cd63-Step1 came from digested products of KpnI/XhoI. The correct plasmid was named Cd63-Step2. The mcherry-bGHpA & phiYFP-rBGpA cassette were cloned into Cd63-Step2 by In-Fusion Enzymes, and the Cd63-Step2 came from digested products of ClaI/AsiSI. The correct plasmid was named Cd63-Step3. The The EGFP-TKpA & mTFP-PGKpA cassette were cloned into Cd63-Step3 by In-Fusion Enzymes, and the Cd63-Step3 came from digested products of AsiSI/XhoI. All steps were subsequently confirmed to be the correct targeting vector by diagnostic PCR, restriction digests and sequencing. This plasmid was subsequently linearized with NotI and used for electroporation to ES cells (C57BL/6).

Negative selection was performed by the expression of diphtheria toxin A (DTA) (present upstream of the HA of the targeting vector), reducing the isolation of non-homologous recombined ES cell clones. Further positive selection with neomycin drug led to 95 drug-resistant clones. PCR screening confirmed 15 potentially targeted clones, 6 of which were validated by Southern Blotting. Next, selected ES cells were used for blastocyst microinjection, followed by chimera production.

### Embryonic stem cells transfection

ESC containing the ExoBow transgene (2×10^5^ cells) were reverse transfected with 2.5μg of each plasmid using LP300. EGFP plasmid was used as a positive control of transfection. After transfection, the mixture was plated in 6 well plates. At the end of the day (after 5h of transfection), the medium was replaced with fresh ES cell medium. After 72h, PCR analysis was performed to evaluate transgene recombination. The following primers were used for the different regions associated with Flp or Cre-mediated recombination:

Region 1 – FLP recombination

Primers for Region 1 (<u>Annealing Temperature 60.0 °C</u>):

> Forward: TGCCTTTTATGGTAATCGTGCGAG
>
> Reverse: CCCACAAAGGCCACCAGGAAGAG

Region 2 – LoxN recombination

Primers for Region 2 (<u>Annealing Temperature 60.0 °C</u>):

> Forward: CTTGCTGCATCAACATAACTGTGG
>
> Reverse: TCCATCTCCACCACGTAGGGGATC

Region 3 – Lox 2272 recombination

Primers for Region 3 (<u>Annealing Temperature 60.0 °C</u>):

> Forward: CTTGCTGCATCAACATAACTGTGG
>
> Reverse: CGTTTACGTCGCCGTCCAGCTCG

Region 4 – Lox5171 recombination

Primers for Region 4 (<u>Annealing Temperature 60.0 °C</u>):

> Forward: CTTGCTGCATCAACATAACTGTGG
>
> Reverse: ATTCACGTTGCCCTCCATCTTCAG

### Mouse lines

KPC ExoBow (*Pdx1-Cre; LSL-Kras*^*G12D/*+^; *LSL-Trp53*^*R172H/*+^; *R26*^*CD63-XFP/*+^) developed spontaneous PDAC tumors in a similar way to the KPC mice. KPC (*Pdx1-Cre*; *LSL- Kras*^*G12D/*+^; *LSL-Trp53*^*R172H/*+^) alleles were purchased from Jackson Laboratory: B6.FVB-Tg(Pdx1-cre)6Tuv/J (IMSR Cat# JAX:014647); B6.129S4-Krastm4Tyj/J (IMSR Cat# JAX:008179); 129S-Trp53tm2Tyj/J (IMSR Cat# JAX:008652).

KPF CD63-mCherry (*Pdx1-Flp; FSF-KRAS*^*G12D/*+^; *Trp53*^*Frt/*+^; *R26*^*CD63-XFP/*+^) developed spontaneous PDAC tumors in a similar way to the KPF mice. All alleles of the KPF mouse model were kindly provided by Dr. Dieter Saur, Technische Universität München, München, Germany.

All mice were housed under standard housing conditions at the i3S animal facility, and all animal procedures were reviewed and approved by the i3S Animal Welfare and Ethics Body and the animal protocol was approved by DGAV “Direção Geral de Alimentação e Veterinária” (ID 015225).

### Polimerase chain reaction

For mice genotyping, an ear fragment was digested at 56°C for 2h on a thermal-shaker with lysis buffer [10 mM Tris (pH 7.5), 400 mM NaCl, 2 mM EDTA (pH 8.0)] (pH 7.3–7.5), 20% sodium dodecyl sulphate (SDS) (Merck 428018), and 20 μL of 20 mg/ml proteinase (Ambion RNA by Life Technologies AM2548)]. Then, 6M NaCl was added to the extraction mixture, samples were mixed thoroughly by vortexing for 10 seconds, followed by centrifugation at 14000RPM for 15 min to precipitate the residual cellular debris. The supernatant was transferred to a clean eppendorf tube and 100% ethanol was added to each sample, mixed thoroughly by vortexing for 10 seconds, and centrifuged at 14000RPM for 5 min to pellet the DNA. The DNA pellets were washed with 70% ethanol, followed by centrifugation at 14000RPM for 5 min. The pellets were completely air dried and resuspended in sterile nuclease-free water. For conventional PCR we used a commercial master mix 2x My Taq HS Mix (Bioline Bio-25046). PCR amplifications were carried out in the T-100 Thermal Cycler (Biorad). All assays included two positive controls samples and a no-template control (contained all reaction components except the genomic DNA). Oligonucleotide sequences used for genotyping were:

R26^CD63-XFP/+^ Forward - CAAAGCTGAAAGCTAAGTCTGCAG

R26^CD63-XFP/+^ Reverse 1 - GGGCCATTTACCGTAAGTTATGTAACG

R26^CD63-XFP/+^ Reverse 2 - GCCATTTAAGCCATGGGAAGTTAG

The amplification protocol included an initial denaturation and enzyme activation at 95°C for 5 min followed by 30 cycles of denaturation at 95°C for 30 seconds, annealing at 55°C for 1 min and 30 seconds, and extension at 72°C for 30 seconds and a final extension at 60°C for 30 seconds.

### *In vivo* imaging system - IVIS

Pancreas were collected and imaged on the IVIS Lumina iii (Caliper) to determine the fluorescence intensities with the appropriate excitation and emission filters. To detect mCherry signal, 535 laser and DsRed emission filter was used, whilst for eGFP, phiYFP and mTFP, 465 laser and GFP emission filter was used. Fluorescence intensity is represented by a multicolor scale ranging from red (least intense) to yellow (most intense). Signal intensity images were superimposed over gray scale reference photographs using IVIS Living image Software.

### Flow cytometry

Pancreas were minced and pieces were meshed through a 70μm strainer (Falcon). The single cell suspension was washed once using Hanks Balanced Salt Solution (HBSS) 1X/FBS 2% and by centrifugation at 600g for 5 min. The pellet was resuspended in HBSS 1X/FBS 2% and filtered through a 35μm cell strainer (Falcon) prior to flow cytometry analysis on BDFACS Aria II Cell Sorter (BD Biosciences). Data from flow cytometry acquisition was analyzed using the FlowJo software (version 10, BD).

### Isolation of extracellular vesicles from pancreas tissue

Pancreas-derived exosomes were isolated according to Crescitelli *et a*.*l* [27]. Briefly, the pancreas were collected and minced into 1-2 mm fragments, followed by 30 min incubation in agitation in RPMI-1640 (Gibco) media supplemented with collagenase D (2 mg/ml, Sigma-Aldrich) and DNase I (40 U/ml, Thermo Fisher Scientific). After enzymatic digestion, samples were meshed through a 70μm strainer (Falcon) using HBBS 1X. Samples were centrifuged at 300g for 10min and 2000g for 20min to remove cells and tissue remains. Cleared supernatants were centrifuged at 16,500 g_avg_ (14,500 rpm) for 20 min and 118,000 g_avg_ (38,800 rpm) for 2.5 h to collect large and small vesicles, respectively.

### Western-blot

Extraction of protein from isolated exosomes was performed using SDS2.5%/8M urea (Sigma-Aldrich) lysis buffer, supplemented with cComplete (Roche) and phenylmethylsulphonyl fluoride (PMSF, Sigma), for 30 min on ice followed by a centrifugation at 17000g for 30 min to remove DNA. 30µg of protein was used for western blot analysis after quantification using DC™ Protein Assay (BIO-RAD). For western-blot analysis of sucrose gradient derived samples, the total volume of 30 μL of each fraction was used. All samples were incubated with laemmli buffer without β-mercaptoethanol (ratio 4:1) for 10 min at 95 °C. Proteins were run in an SDS-PAGE (sodium dodecylsulphate-polyacrylamide gel electrophoresis) gel. The Precision Plus Protein™ Dual Color Standards (BIO-RAD) was used as ladder control. After separation by electrophoresis, proteins were transferred onto nitrocellulose membranes 0.2μm (GE Healthcare) using a wet electrophoretic transfer. Ponceau S staining was used to confirm an effective and equilibrated protein transfer. Subsequently, the nitrocellulose membranes were blocked with 5% non-fat dry milk in phosphate-buffered saline (PBS) 1X/0.1% Tween 20 (Sigma-Aldrich) for 1 hour at room temperature. After blocking, membranes were incubated overnight at 4°C on a shaker with the following primary antibodies: anti-mCherry 1:500, Biorbyt orb116118, anti-eGFP 1:500, Abcam ab13970, anti-phiYFP 1:1000, Evrogen AB603 and anti-mTFP 1:500, kindly provided by Cai Laboratory, University of Michigan Medical School, Michigan, USA. After 4 washes with PBS 1X/ 0.1% Tween 20, membranes were incubated 1 hour at room temperature with horseradish peroxidase (HRP)-conjugated secondary antibodies: anti-goat 1:5000 Abcam ab6741, anti-chicken 1:5000 Sigma-Aldrich A906, anti-rabbit 1:5000 Cell Signalling 7074 and anti-rat 1:5000, GenScript a00167, respectively. Membranes were washed with PBS 1X/0.1% Tween 20 and incubated with Clarity™ Western Enhanced chemiluminescence (ECL) Substrate (BIO-RAD), according to the manufacturer’s recommendations, to detect the antibody-specific signal using GE Healthcare Amersham™ Hyperfilm™ ECL.

### Immunofluorescence

#### Cell lines

BxPC-3 or KPC cells were plated in coverslips and 24 hours after media was removed and cells were washed with cold PBS 1X followed by fixation with 4% paraformaldehyde (PFA) (Sigma-Aldrich) for 15 min at room temperature. Coverslips were rinsed three times with PBS1X and incubated with a quenching solution of 0.1M glycine for 5 min at room temperature. The cells were permeabilized with a solution of Triton-X (VWR) 0.1% followed by a 45 min incubation at room temperature with 10% Albumine Bovine Fraction V (BSA) (NZYTech). After blocking, cells were incubated overnight at 4°C with the following primary antibodies: anti-human CD63 (Novus Biologicals® H5C6), anti-eGFP (BIO-RAD 81/4745-1051), anti-mCherry, anti-phiYFP and anti-mTFP, in a dilution of 1:500 in PBS 1X/ 2% BSA. Anti-mCherry, anti-phiYFP and anti-mTFP antibodies were developed and kindly provided by Cai Laboratory, University of Michigan Medical School, Michigan, USA. The KPC cell line was only incubated with the anti-human CD63 antibody. Next day, after 4 washes with PBS 1X, cells were incubated 45 min at room temperature with the respective secondary antibodies: anti-mouse Alexa-Fluor® 594 (abcam, ab150108), anti-chicken Alexa-Fluor® 633 (Sigma, SAB4600127), anti-sheep Alexa-Fluor® 488 (Jackson Immunoresearch, 713-545-003), anti-rabbit Alexa-Fluor® 488 (Jackson Immunoresearch, 711-545-152) and anti-rat Alexa-Fluor® 488 (Invitrogen, A21208), at a 1:500 dilution in PBS 1X/2% BSA. Nuclei counterstain was achieved using Hoechst solution (1:10 000, Thermo Scientific) for 10 min at room temperature. The coverslips were mounted in glass slides using a drop of VECTASHIELD mounting medium (Vector Laboratories) and sealed with nail polish.

### Formalin-fixed organs

Tissue samples were fixed in 10% formalin for at least 24 hours prior to paraffin embedding. Next, 6μm thick sections were cut using a Microm HM335E microtome, transferred to coated slides (Thermo Fisher Scientific), and left overnight at 37 °C. Sections were deparaffinized and hydrated prior to heat-mediated antigen retrieval with sodium citrate buffer pH 6.0 (Vector Laboratories) for 40 min inside a steamer machine. Afterwards, incubation with a 0.3% hydrogen peroxide in methanol solution (H^2^O^2^ [Sigma-Aldrich]; CH^3^OH [VWR]) for 15 min at room temperature. After washes with PBS 1X/0.1% Tween 20 tissue sections were bordered with a hydrophobic pen (Vector Laboratories), placed in the humid chamber and incubated with Ultravision Protein-block solution (DAKO) for 1 hour at room temperature. Incubation with primary antibody anti-mCherry 1:500, anti-α-SMA (1:400 Sigma-Aldrich, a2547) was performed at 4°C for one and a half days in PBS 1X/0.1% Tween 20. After primary antibody incubation, slides were washed three times with PBS 1X/0.1% Tween 20 for 10 min. In the humid chamber, the secondary antibody incubation with anti-rabbit Alexa-Fluor® 488 1:500 (Jackson Immunno Research Lab, JK711545152), and anti-mouse Alexa-Fluor® 594 1:500 (abcam, ab150108), respectively, was performed. After incubation, the slides were thoroughly washed with PBS 1X/0.1% Tween 20. The nuclei were counterstained with Hoechst solution (1:10 000, Thermo Scientific) for 15 min at room temperature. The slides were washed and mounted with VECTASHIELD mounting medium (Vector Laboratories) and sealed with nail polish.

### Paraformaldehyde-fixed organs

Mice were anesthetized with ketamine and medetomidine and transcardially perfused using ice-cold PBS 1X followed by ice-cold 4% paraformaldehyde (PFA) in PBS 1X. Pancreas were collected and post-fixed overnight in 4% PFA, rinsed and incubated overnight in 30% (weight/volume) sucrose solution. After that, organs were embedded in O.C.T. and frozen using dry ice. Frozen sections of about 15μm were made using the Cryostat Leica CM 3050S (Leica Biosystems). Sections were dried for 1 hour at room temperature followed by washes with PBS 1X. Sodium citrate-based antigen retrieval solution pH 8.5 was pre-heated in a water bath at 80°C, and slides incubated for 30 min. Slides were rinsed using PBS 1X/Triton-X 0.1% and incubated overnight at 37°C in PBS 1X/Triton-X 1% with gentle shaking for permeabilization, followed by 10 hours incubation in blocking solution (PBS 1X/10% BSA) supplemented with 1% Triton-X at 4°C with gentle shaking. Next, tissue sections were delimited with a hydrophobic pen (Vector Laboratories), placed in the humid chamber and incubated at 4°C for one and a half days with the primary antibodies prepared in PBS 1X/Triton-X 0.5%: anti-α-SMA (1:400 Sigma-Aldrich, a2547), anti-GFP 1:300 (BIO-RAD 4745-1051), anti-mCherry 1:500, anti- phiYFP 1:300, anti-mTFP 1:500. Anti-reporter antibodies were developed and kindly provided by Cai Laboratory, University of Michigan Medical School, Michigan, USA. After incubation with primary antibodies, slides were thoroughly washed using PBS 1X/Triton-X 0.1% and incubated overnight at 4°C with the respective antibodies anti-mouse Alexa-Fluor® 594 (abcam, ab150108), anti-sheep Alexa-Fluor® 488 (Jackson Immunoresearch, 713-545-003), anti-rabbit Alexa-Fluor® 488 (Jackson Immunno Research Lab JK711545152), anti-rabbit Alexa-Fluor® 546 (ThermoFisher Scientific, A10040) and anti-rat Alexa-Fluor® 594 (ThermoFisher Scientific, A21209), in a dilution 1:500. Slides were thoroughly washed and the nuclei counterstained with Hoechst solution (1:10 000, Thermo Scientific) for 15 min at room temperature. The slides were washed and mounted with VECTASHIELD mounting medium (Vector Laboratories) and sealed with nail polish.

### Microscopy Imaging

A Leica TCS SP5 inverted confocal system was used to image samples with an HCL PL APO CS 63x/NA 1.40 oil objective or 20x/0.70 IMM/CORR water objective, the later in Figure 4D. Sequential acquisition was used to acquire each fluorescent signal apart from 405 laser for Hoescht detection. An upright Zeiss LSM 780 was also used in Figure 4E and 5C, with 40x 1.3NA oil immersion objective. Presented images were max z-projected (when referenced in figure legend), cropped and contrast was optimized in FIJI.

## Acknowledgments

The work was supported by the project NORTE-01-0145-FEDER-000029, Norte Portugal Regional Programme (NORTE 2020), under the PORTUGAL 2020 Partnership Agreement, through the European Regional Development Fund (ERDF) and national funds through FCT—Foundation for Science and Technology POCI-01-0145-FEDER- 32189. BA, NB and CFR are supported by FCT (PD/BD/135546/2018; SFRH/BD/130801/2017 and SFRH/BD/131461/2017). The authors acknowledge the support of the i3S Scientific Platforms: Translational Cytometry, Animal Facility, Bioimaging and Histology and Electron Microscopy (HEMS). Bioimaging and HEMS are members of the national infrastructure PPBI - Portuguese Platform of Bioimaging (PPBI-POCI-01-0145-FEDER-022122).

## Author contributions

Conceptualization, S.A.M; B.A. and C.F.R; Methodology, B.A., D.C. and S.A.M.; Formal analysis, B.A.; Investigation, B.A., N.B., C.F.R. and P.F.V.; Resources, D.S., B.S., D.C. and S.A.M.; Data curation: B.A.; Writing - original draft, B.A. and S.A.M; Writing – Review & Editing, N.B., J.C.M, and S.A.M; Visualization, B.A.; Supervision, S.A.M. Funding acquisition, S.A.M.

## Figures legend

**Supplementary Figure 1 (related to Figure 1).**
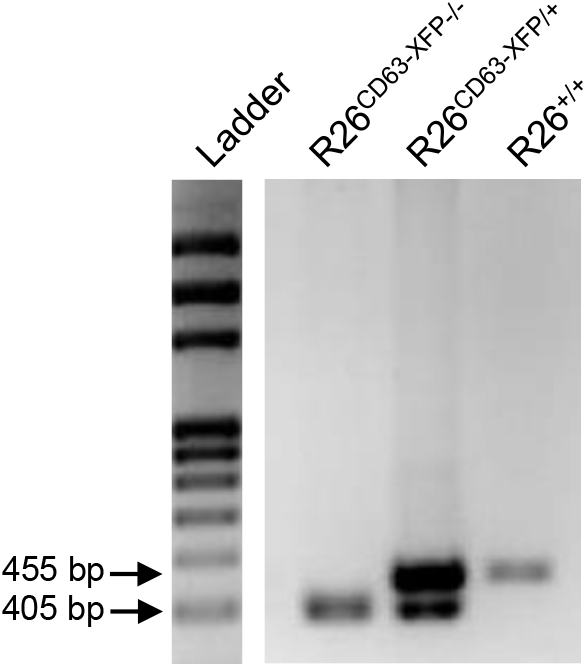
Representative genotyping PCR of the ExoBow transgene (R26^CD63-XFP^) in DNA derived from the ear of a R26^CD63-XFP-/-^homozygote mouse, R26^CD63-XFP/+^heterozygous mouse and WT mouse (R26^+/+^). R26 WT allele 455 bp, R26^CD63-XFP^allele 405 bp.

**Supplementary Figure 2 (related to Figure 3).**
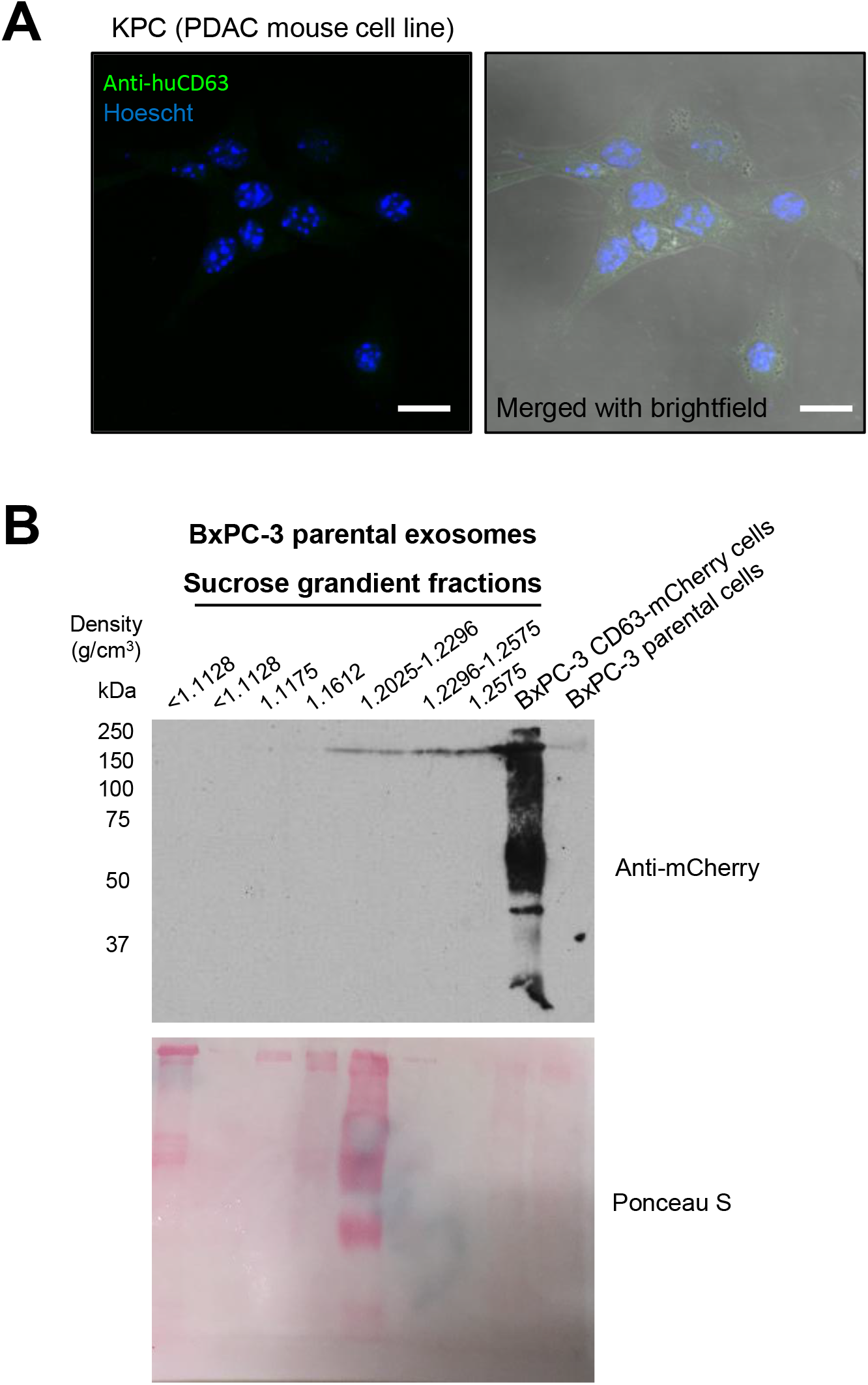
**(A)** Confocal microscopy image of a maximum projection of KPC parental mouse cells immunostained against anti-human CD63 (green). The nucleus counterstained with Hoescht are represented in blue. Merged confocal microscopy image with brightfield (right panel). Scale bar 20μm. Negative control to demonstrate antibody specificity to human CD63. **(B)** Representative western-blot of CD63-mCherry expression in cells and exosomes derived from the BxPC-3 parental cell line. Individual 1 mL fractions were collected and after ultracentrifugation were loaded on gels for electrophoresis. Exosomes are located in fractions around 1.1415 to 1.2025 g/cm^3^density. BxPC-3 CD63-mCherry and parental cells were used as positive and negative controls, respectively.

**Supplementary Figure 3 (related to Figure 4).**
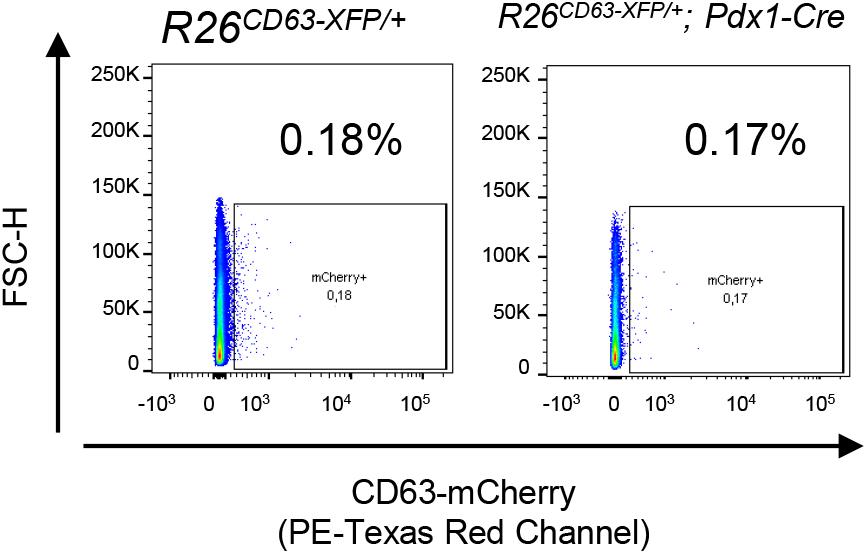
Representative flow cytometry analysis of CD63-mCherry in the pancreas of R26^CD63-XFP/+^and *Pdx1-Cre R26*^*CD63-XFP/*+^mice. These correspond to negative controls to demonstrate that ExoBow transgene expression is dependent on flippase-mediated recombination.

## Full western-blots

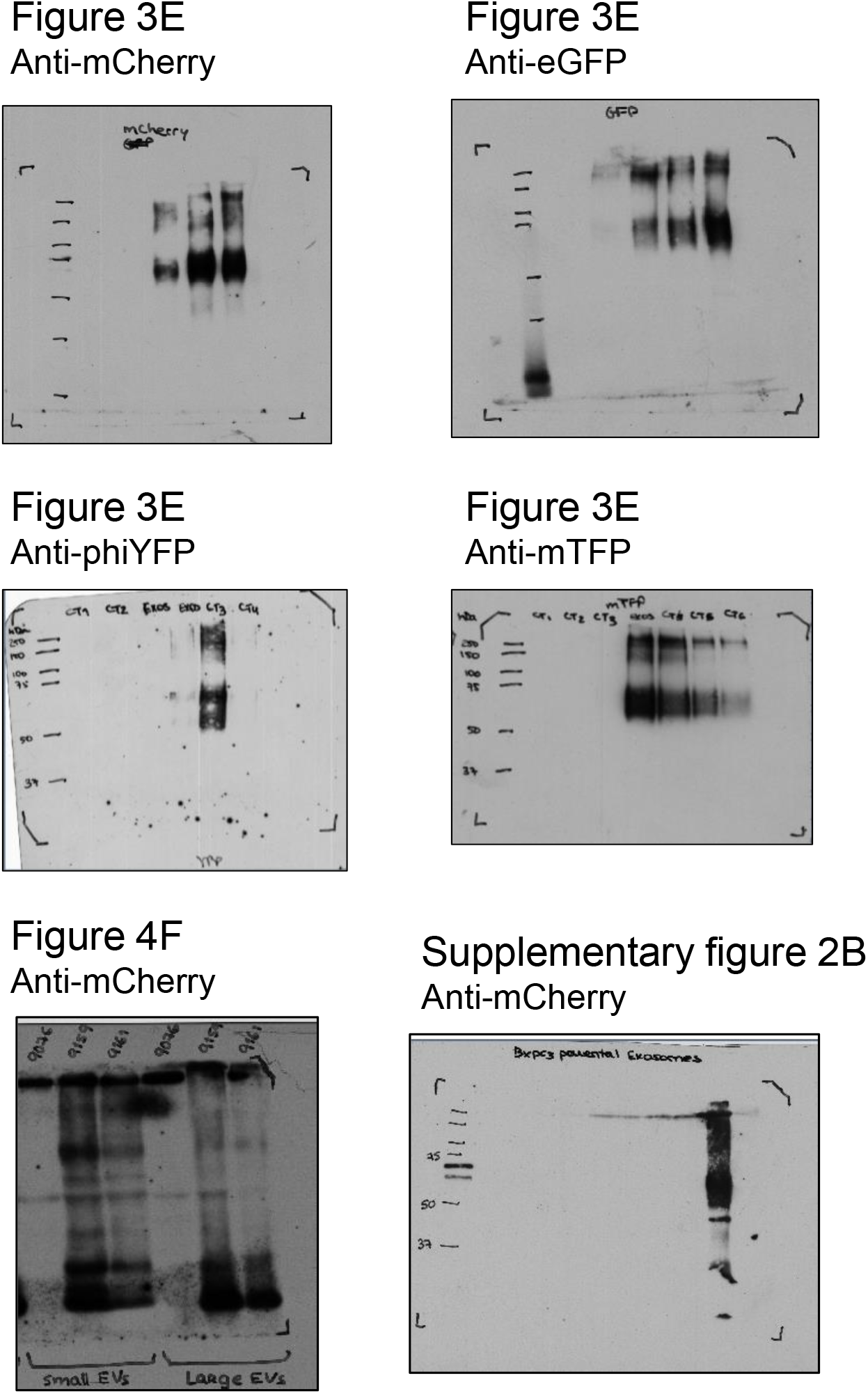

